# Towards a new kind of vaccine

**DOI:** 10.1101/101345

**Authors:** Reginald M. Gorczynski, Geoffrey W. Hoffmann

## Abstract

We present new data showing that normal IgG immune responses comprise the production of two kinds of antibodies, namely anti-foreign and anti-anti-self antibodies. For example, immunization of C3H mice by two rounds of BL/6 skin grafting results in the production of anti-BL/6 antibodies plus antiidiotypic antibodies (C3H anti-anti-C3H) with the latter being detected using antibodies produced in a BL/6 anti-C3H immune response. Similarly, the IgG immune response of C3H mice to tetanus toxoid includes the production of C3H anti-anti-C3H antibodies. Antigen-specific antibodies produced in one alloimmunization plus antiidiotypic antibodies produced in the converse immunization can be used to synergistically induce specific tolerance. We show that infusions of anti-BL/6 antibodies together with BL/6 anti-anti-BL/6 antibodies specifically suppress an immune response to BL/6 lymphocytes in C3H mice. Specific tolerance was measured as suppression of the induction of BL/6-specific cytotoxic T cells. The two kinds of antibodies with complementary specificity are believed to stimulate two populations of T lymphocytes, and co-selection (mutual selection) of these two populations leads to a new stable steady state of the system that has specifically diminished reactivity to BL/6 tissue. Stimulation with a combination of anti-C3H and C3H anti-anti-C3H IgG antibodies furthermore down-regulates inflammation in a mouse model of inflammatory bowel disease. An analogous combination of C3H anti-BL/6 and BL/6 anti-anti-BL/6 antibodies significantly down-regulates tumour growth and metastases in BALB/c mice in the EMT6 transplantable breast cancer model. We conclude that a combination of certain antigen-specific and antiidiotypic antibodies has potential as a new class of vaccines based on the symmetrical immune network theory. This new kind of vaccine does not involve the production of antibodies. The prevention of two important degenerative diseases makes this a potential anti-aging technology.

## Introduction

Much research on antiidiotypic antibodies has been focused on such antibodies mimicking the shape of the antigen, and therefore being able to substitute for the antigen (Charmat et al., 1984, Guillet et al., 1985, Urbain et al., 1979) An antibody specific for an antigen X has a V region that is anti-X, and we say it has an anti-X idiotype. If we immunize a rabbit with an anti-X antibody together with an adjuvant the rabbit may make anti-anti-X antiidiotypic antibodies. However, such antiidiotypic antibodies play no role in the immune network theory that has been developed in a series of papers and a monograph published from 1975 to the present (Hoffmann et al., 1986; Hoffmann, 1994; Hoffmann, 2008; Leung and Hoffmann, 2014). The theory is called the symmetrical immune network theory. The antiidiotypes that play a role in the symmetrical immune network theory are of two types, namely second symmetry antiidiotypes (Hoffmann et al., 1986) and co-selection antiidiotypes (Hoffmann, 1994). Second symmetry antiidiotypes are for example A anti-anti-A antibodies that are present in an A anti-B serum, that are specific for B anti-A antibodies, that in turn are present in a B anti-A immune response. Co-selection antiidiotypes are expressed by lymphocytes that are co-selected (mutually selected) with antigen-specific lymphocytes.

In this paper, we firstly extend the ways in which second symmetry antiidiotypes can be produced. In addition to previously demonstrated immunization with lymphocytes, immunization with skin grafts and with a protein antigen plus an adjuvant result in the production of second symmetry antiidiotypes.

We then show that a mutually specific pair of antibodies such as anti-A and anti-anti-A synergistically induce tolerance in a vertebrate C. Furthermore, infusions with such a combination of antibodies strengthens the immune system of C such that C becomes resistant to both inflammatory bowel disease and a transplantable breast cancer. Two immune systems, A and B, are typically involved in producing the anti-A and the anti-anti-A antibodies, and the immune system of the treated vertebrate C is also involved in making C’s immune system stronger. C may be the same as or different from B. Hence the strengthened immune system of C is the result of the participation of two or three immune systems working together.

## Results

### Second symmetry antiidiotypic antibodies induced by skin grafting and immunization with a protein antigen

It has previously been shown that immunization of a strain A mouse with lymphocytes of a strain B can induce the production of both antigen-specific (A anti-B) and antiidiotypic (A anti-anti-A) antibodies (Hoffmann et al., 1986). Conversely, immunization of B with A lymphocytes induces B anti-A and B anti-anti-B antibodies.

The antigen-specific antibodies in the A anti-B serum are specific for the antiidiotypic antibodies (B anti-anti-B) in the B anti-A serum. Similarly, the antigen-specific antibodies in a B anti-A serum are specific for the antiidiotypic antibodies (A anti-anti-A) in the A anti-B serum. This set of relationships is known as “second symmetry” (Hoffmann, 2008, chapter 14). ("First symmetry" in the symmetrical immune network theory is the concept that if an idiotype A is antiidiotypic to an idiotype B, the idiotype B is antiidiotypic to the idiotype A (Hoffmann, 2008, chapter 9).

In our previous work on second symmetry in mice we generated A anti-anti-A antibodies (antiidiotypic antibodies) by immunizing mice of strain A with strain B lymphocytes. The A anti-anti-A antibodies were assumed to be produced in response to anti-A receptors of the strain B lymphocytes. We have however now found that we do not need to immunize strain A with lymphocytes to generate the A anti-anti-A immune response. Rather, two rounds of B strain skin grafts on A strain mice also induce the production of A anti-anti-A antibodies in the A strain mice. This is shown in the ELISA assays of Figures 1A and 1B.

**Figure 1.**
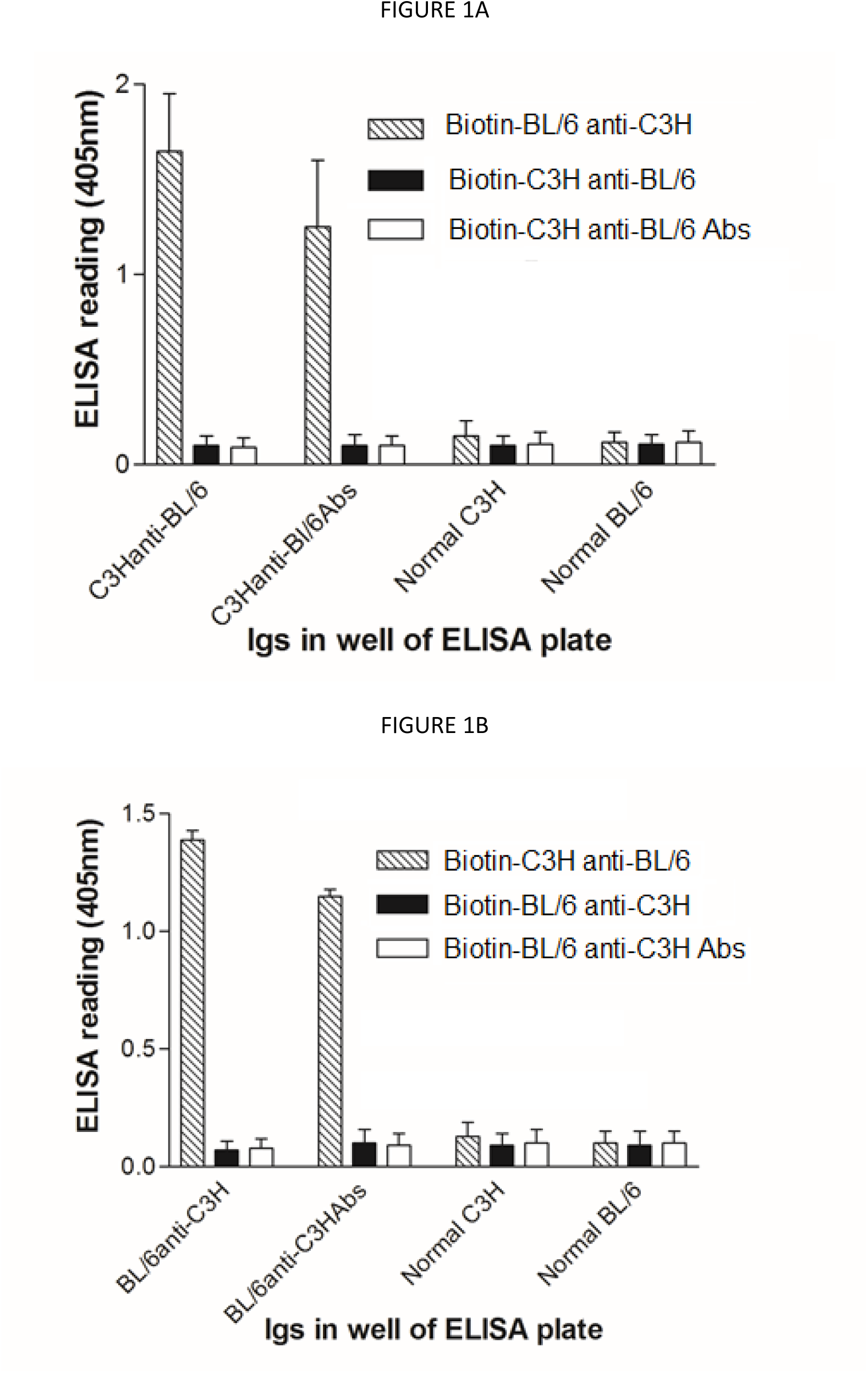
Production of antiidiotypic antibodies in IgG immune responses to skin grafts or to a protein antigen. Figure 1A (top) shows biotinylated BL/6 anti-C3H IgG binds to C3H anti-BL/6 IgG and to C3H anti-BL/6 IgG absorbed with BL/6 lymphocytes, but not to normal C3H IgG, nor to normal BL/6 IgG. Additional controls are biotinylated C3H anti-BL/6 IgG and biotinylated (C3H anti-BL/6 IgG absorbed with BL/6 lymphocytes). Conversely, Figure 1B (bottom) shows biotinylated C3H anti-BL/6 IgG binds to BL/6 anti-C3H IgG and to BL/6 anti-C3H IgG absorbed with C3H lymphocytes, but not to normal BL/6 IgG, nor to normal C3H IgG. Additional controls are biotinylated BL/6 anti-C3H IgG and biotinylated (BL/6 anti-C3H absorbed with C3H lymphocytes). In all cases 10ng IgG was coated to wells of the ELISA plate, and biotinylated IgGs for detection were used at 1:2500 concentration (with streptavidin-HRP at 1:2000).

In the ELISA assays, normal IgG, alloimmune IgG or IgG from alloimmune mice that had been absorbed using lymphocytes of the strain used in the skin grafting, were coated on ELISA plates at the dilutions shown in the caption of Figure 1. After incubation with biotinylated IgG purified from allo-antiserum, or biotinylated control IgG, the bound antibodies were detected with strep avidin-alkaline phosphatase and substrate. The resulting signal is interpreted as resulting from the binding of biotinylated antigen-specific antibodies to complementary antiidiotypic antibodies on the plate. Figure 1A shows biotinylated BL/6 anti-C3H IgG binds to C3H anti-BL/6 IgG and to C3H anti-BL/6 IgG absorbed with BL/6 lymphocytes, but not to normal C3H IgG, nor to normal BL/6 IgG. This is ascribed to the presence of C3H anti-anti-C3H antibodies in the C3H anti-BL/6 IgG and in the C3H anti-BL/6 IgG absorbed with BL/6 lymphocytes. Absorption with BL/6 lymphocytes removes antigen-specific (anti-BL/6) antibodies without removing the antiidiotypic antibodies. Additional controls are biotinylated C3H anti-BL/6 IgG and biotinylated (C3H anti-BL/6 IgG absorbed with BL/6 lymphocytes). Conversely, Figure 1B shows biotinylated C3H anti-BL/6 IgG binds to BL/6 anti-C3H IgG and to BL/6 anti-C3H IgG absorbed with C3H lymphocytes, but not to normal BL/6 IgG, nor to normal C3H IgG. Additional controls are biotinylated BL/6 anti-C3H IgG and biotinylated (BL/6 anti-C3H IgG absorbed with C3H lymphocytes) (P < 0.01 by ANOVA).

In Table 1 we see the set of interactions between antibodies 1 on the plate (Antibody 1), and the biotinylated antibodies (Antibody 2).

**TABLE 1.**
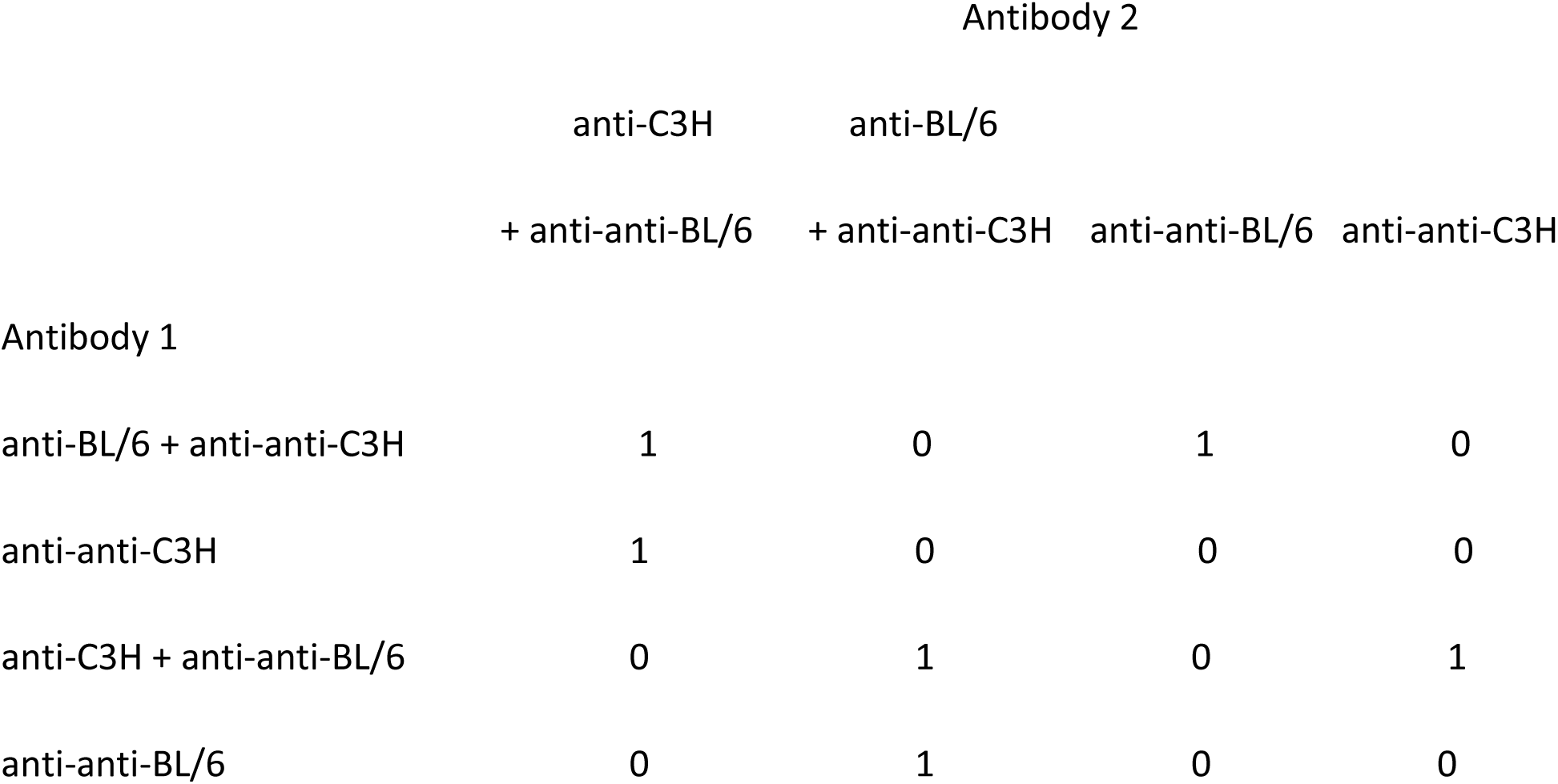

A strong ELISA signal is denoted by 1, a much weaker (background) signal is entered as a 0 in this table. Anti-C3H IgG contained in (anti-C3H + anti-anti-BL/6) IgG plus anti-anti-C3H IgG define an anti-C3H-anti-anti-C3H shape space axis that is orthogonal to the anti-BL/6-anti-anti-BL/6 shape space axis defined by anti-BL/6 IgG contained in (anti-BL/6 + anti-anti-C3H) IgG plus anti-anti-BL/4 IgG.

We have furthermore found that an IgG response to a protein antigen given in an adjuvant induces both an antigen-specific response and an antiidiotypic response. We measured the production of antiidiotypic antibodies in BL/6 and C3H mice immunized with tetanus toxoid (Td) in the adjuvant MPLA. Six BL/6 mice and six C3H mice were immunized on day 0 and day 14 and bled on day 21. Normal IgG, IgG purified from the immunized mice, or IgG from the immunized mice that had been absorbed with Td were coated on ELISA plates at the dilutions shown in Figures 1C and 1D. Figure 1C shows that biotinylated BL/6 anti-C3H IgG binds to C3H anti-Td IgG and to C3H anti-Td IgG absorbed with Td, but not to normal C3H IgG, nor to normal BL/6 IgG. This is ascribed to the presence of C3H anti-anti-C3H antibodies in the C3H anti-Td IgG and in the C3H anti-Td IgG absorbed with Td. Additional controls are biotinylated C3H anti-BL/6 IgG and biotinylated (C3H anti-BL/6 IgG absorbed with BL/6 lymphocytes). Conversely, Figure 1D shows that biotinylated C3H anti-BL/6 IgG binds to BL/6 anti-Td IgG and to BL/6 anti-Td IgG absorbed with Td, but not to normal BL/6 IgG, nor to normal C3H IgG. Additional controls are biotinylated BL/6 anti-C3H IgG and biotinylated (BL/6 anti-C3H IgG absorbed with C3H lymphocytes) (P < 0.05 by ANOVA). This confirms that the IgG immune response of C3H and of BL/6 mice to Td includes the production of C3H anti-anti-C3H and BL/6 anti-anti-BL/6 antibodies respectively.

**Figure 1C.**
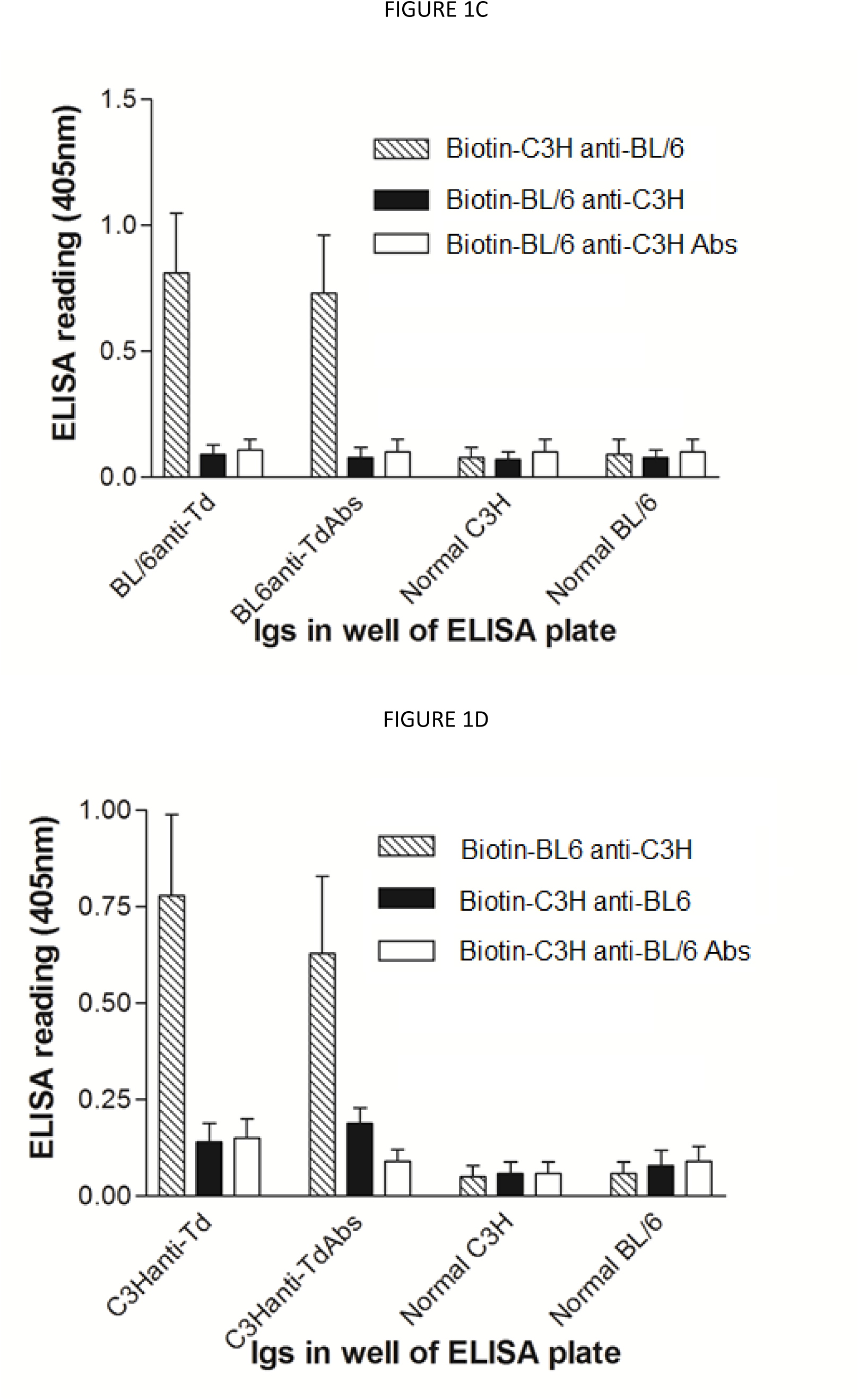
Production of anti-foreign and anti-anti-self antibodies in BL/6 and C3H mice immunized with tetanus toxoid (Td) in the adjuvant MPLA. Normal IgG, IgG purified from the immunized mice, or IgG from the immunized mice that had been absorbed with Td were coated on ELISA plates at 50ng/well. Test developing sera were used at 1:400 concentration, with streptavidin HRP used at 1:2000. Figure 1C (top) shows that biotinylated BL/6 anti-C3H IgG binds to C3H anti-Td IgG and to C3H anti-Td IgG absorbed with Td, but not to normal C3H IgG, nor to normal BL/6 IgG. Additional controls are C3H anti-Td and BL/6 anti-Td absorbed with Td. Conversely, Figure 1D (bottom) shows that biotinylated C3H anti-BL/6 IgG binds to BL/6 anti-Td IgG and to BL/6 anti-Td IgG absorbed with Td, but not to normal BL/6 IgG, nor to normal C3H IgG. Additional controls are C3H anti-Td and C3H anti-Td absorbed with Td. Thus the IgG response to Td includes the production of strain-specific anti-anti-self antibodies.

### Synergy between antigen-specific plus antiidiotypic antibodies in inducing tolerance

Specific tolerance can be induced using co-selection and the above second symmetry relationships. We postulated that a vertebrate A can be treated with a combination of A anti-B antibodies (antigen-specific) and B anti-anti-B antibodies (antiidiotypic) to induce a state in which there is tolerance in A that is specific for B. The envisaged mechanism is shown in Figure 2. Figure 2A shows how the A anti-B antibodies stimulate anti-anti-B T cells, and the anti-anti-B antibodies stimulate anti-B T cells in a vertebrate with an A or C phenotype. There is co-selection of the anti-anti-B T cells and the anti-B T cells, taking the immune system of A or C to a state in which there are elevated levels of these two T cell populations. This is a state in which A or C is specifically unresponsive to B. Figure 2B shows an example in which A is C3H strain mice, B is BL/6 mice, and C is BALB/c mice. The B anti-anti-B antibodies are produced by absorbing the IgG from the hyperimmunized B strain animals with A strain lymphocytes to remove B anti-A IgG. Purified IgG from the converse hyperimmune serum is used without absorption. An A strain vertebrate or a vertebrate of the third phenotype C receives infusions of the A anti-B plus B anti-anti-B IgG. In the latter case, using the third phenotype vertebrate C, the treatment induces a new stable steady state with participation of the immune systems of three vertebrates A, B and C. In the application to a treatment for people, A, B may be two strains of rodents, and C will be a treated person.

**Figure 2.**
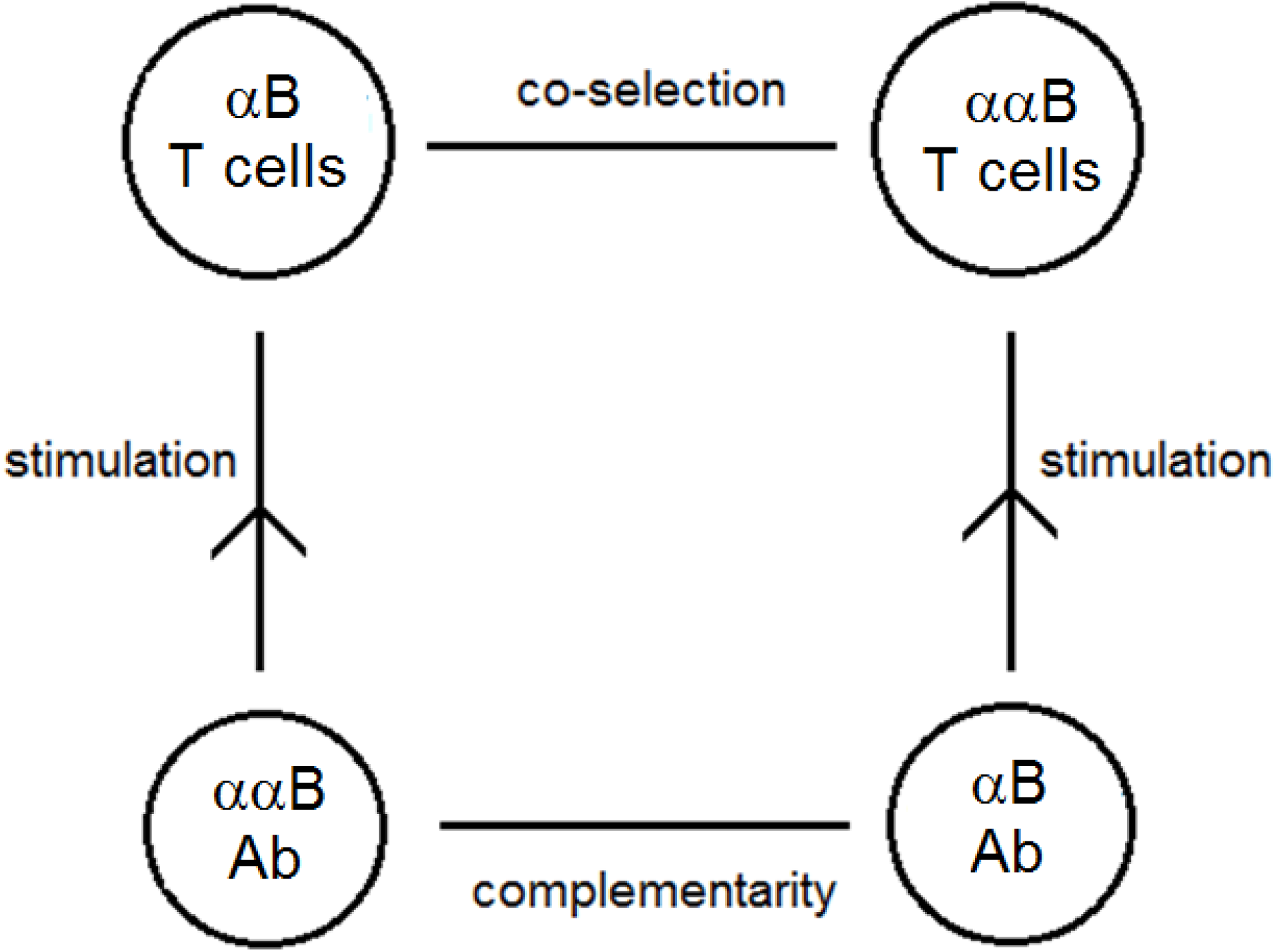
Co-selection mechanism for inducing transplantation tolerance in a vertebrate C using anti-B plus anti-anti-B antibodies. **Figure 2A.** The Greek letter α is an abbreviation for “anti-”. Anti-B antibodies stimulate anti-anti-B T cells and the anti-anti-B antibodies stimulate anti-B T cells. There is co-selection of the anti-anti-B T cells and the anti-B T cells, taking the immune system of C to a state in which there are elevated levels of these two T cell populations. This is a state of the vertebrate C that is specifically suppressed with respect to making an immune response to B.

**Figure 2B.**
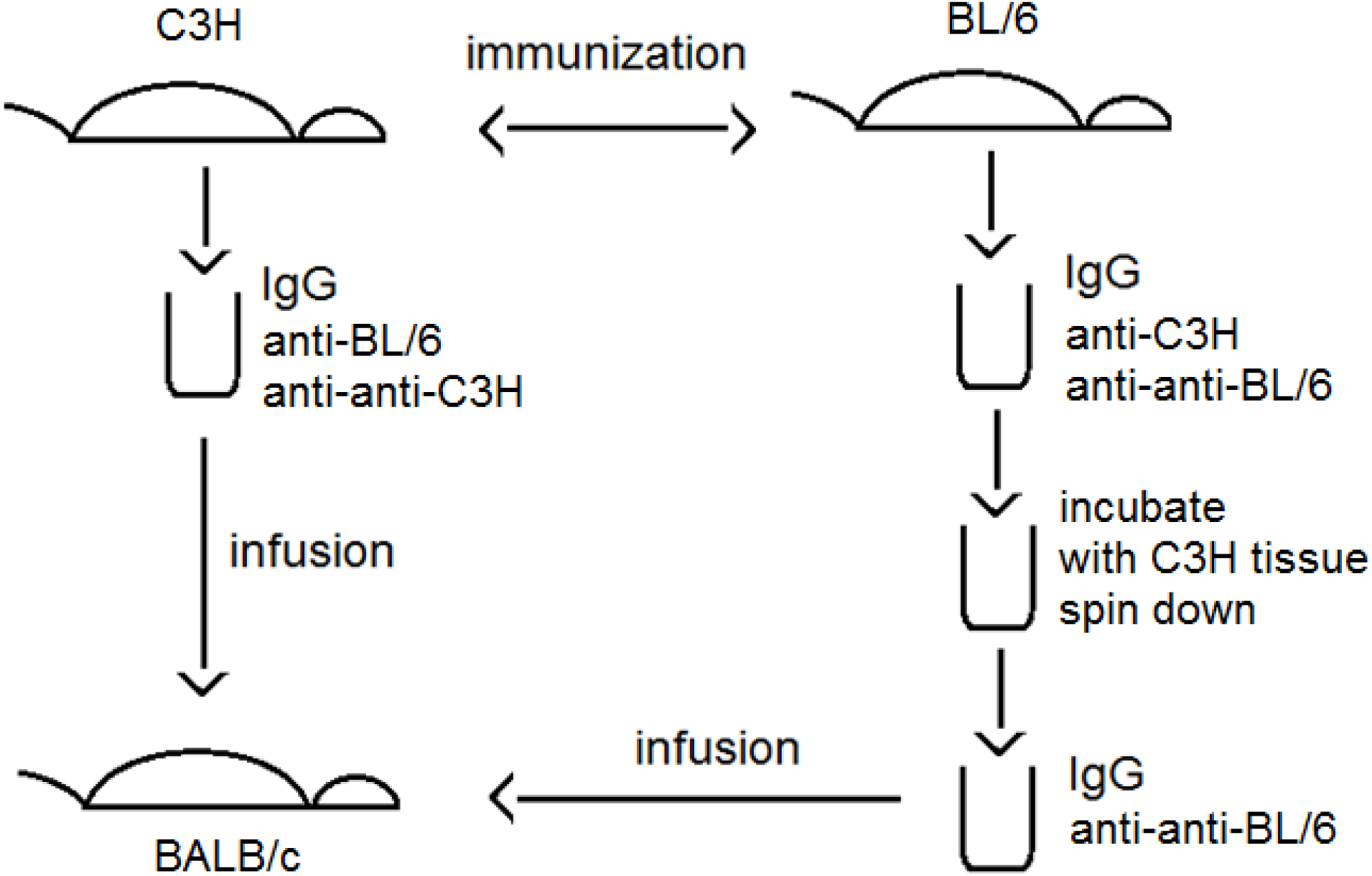
Example showing how three immune systems (BL/6, C3H and BALB/c) are combined to make a stronger than normal immune system in the treated vertebrate (BALB/c). Immunization of strain BL/6 with strain C3H tissue results in the production of anti-C3H plus anti-anti-BL/6 IgG, and the converse immunization results in the production of anti-BL/6 plus anti-anti-C3H IgG. The anti-C3H antibodies in the former IgG are removed by absorption with C3H tissue, for example absorption with C3H lymphocytes. Spinning the preparation down yields a supernatant enriched in anti-anti-BL/6 IgG and without anti-C3H antibodies. A vertebrate of a third phenotype, here BALB/c, receives infusions of anti-anti-BL/6 plus anti-BL/6 IgG. The BALB/c mice also receive anti-anti-C3H IgG, but this is irrelevant because they receive no anti-C3H IgG, and hence there is no C3H-specific positive feedback loop.

We have tested this concept in an experiment in which A was C3H mice and B was BL/6 mice. The C3H anti-BL/6 IgG antibodies were contained in serum obtained from C3H mice that had undergone two rounds of skin grafting with BL/6 skin. The BL/6 anti-anti-BL/6 antibodies were obtained from BL/6 mice that had undergone two rounds of skin grafting with C3H skin. The IgG from the latter mice was absorbed with C3H spleen cells and thymocytes until all detectable anti-C3H antibodies had been removed. The results of two experiments are shown in Figure 3.

**Figure 3.**
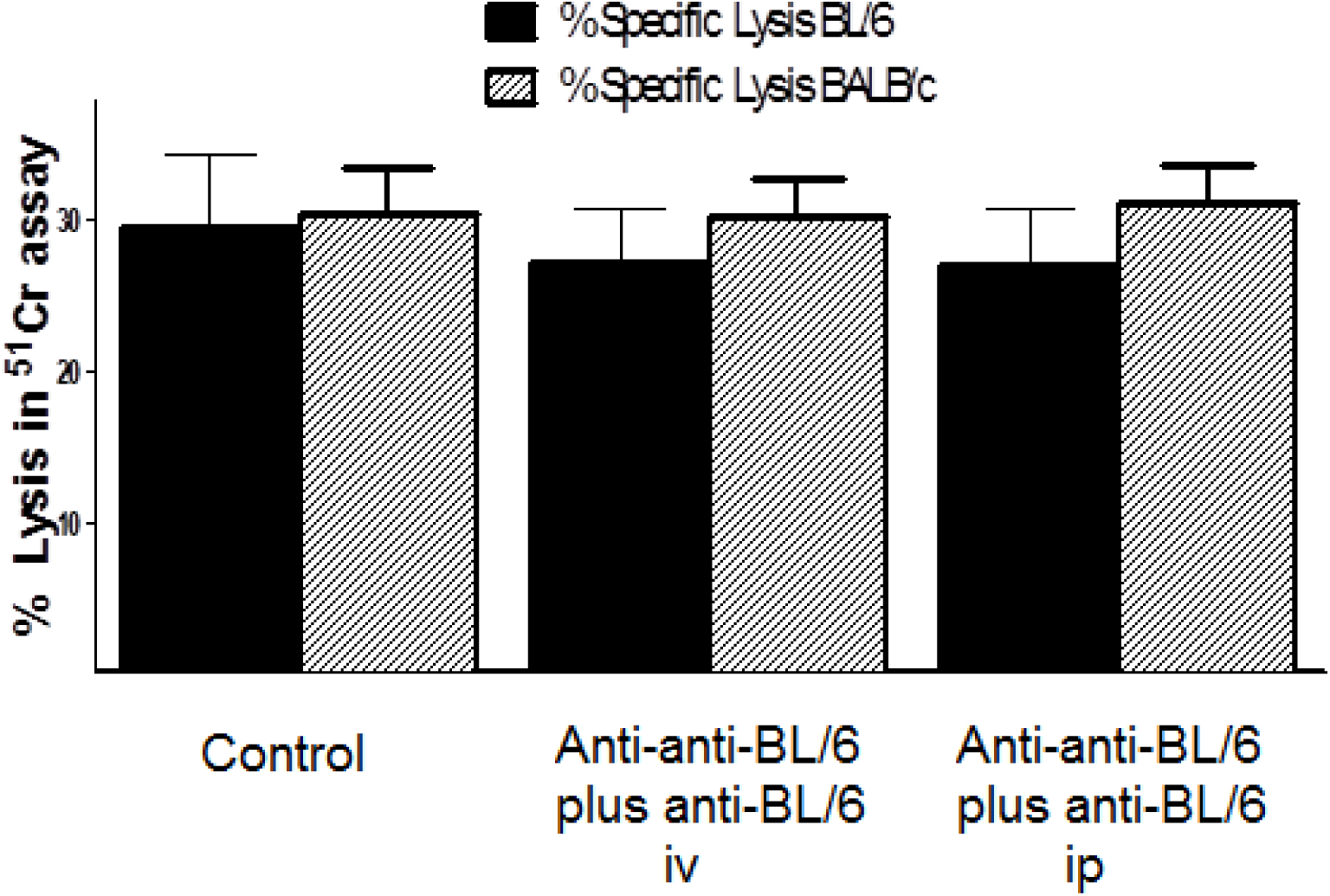
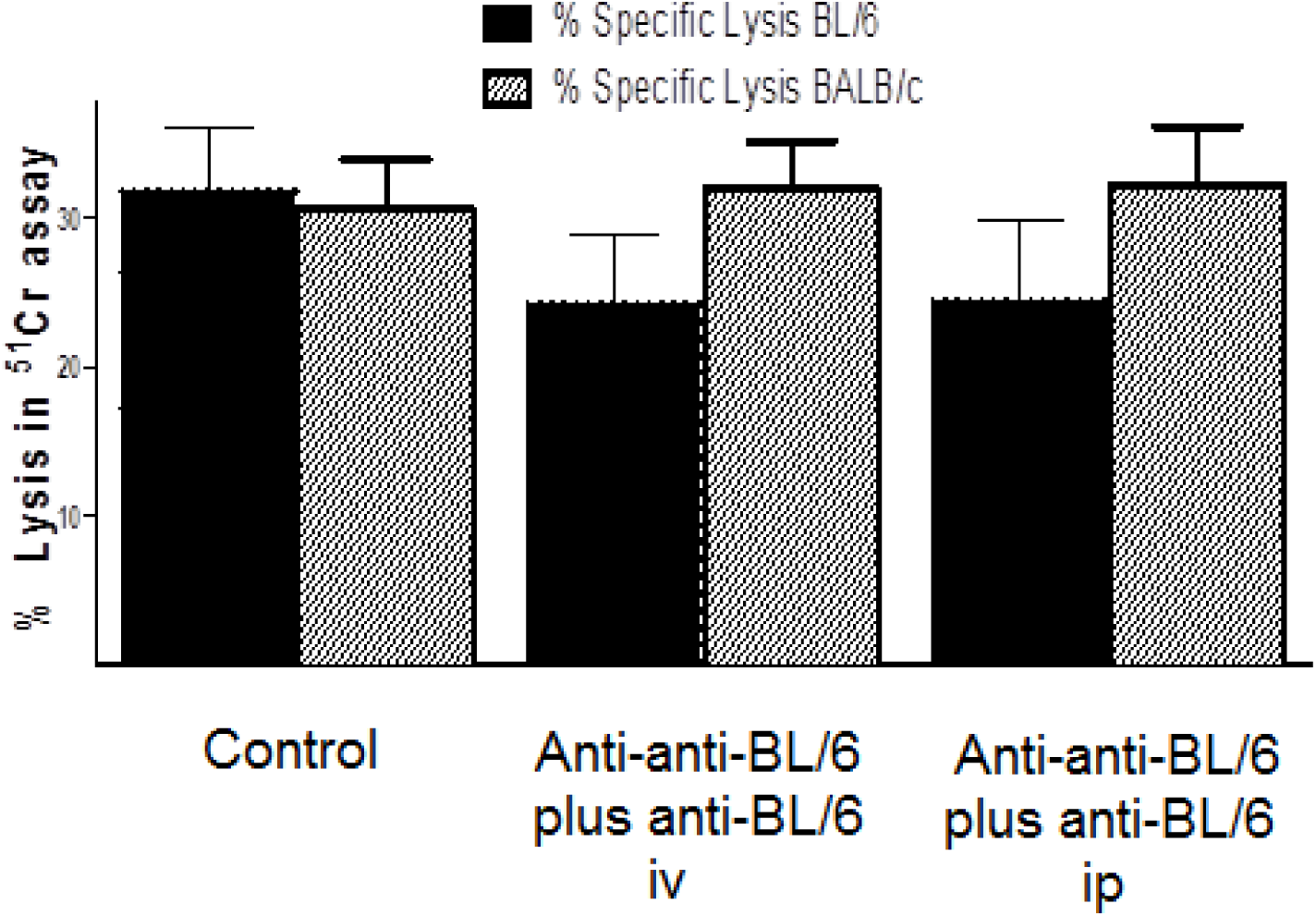
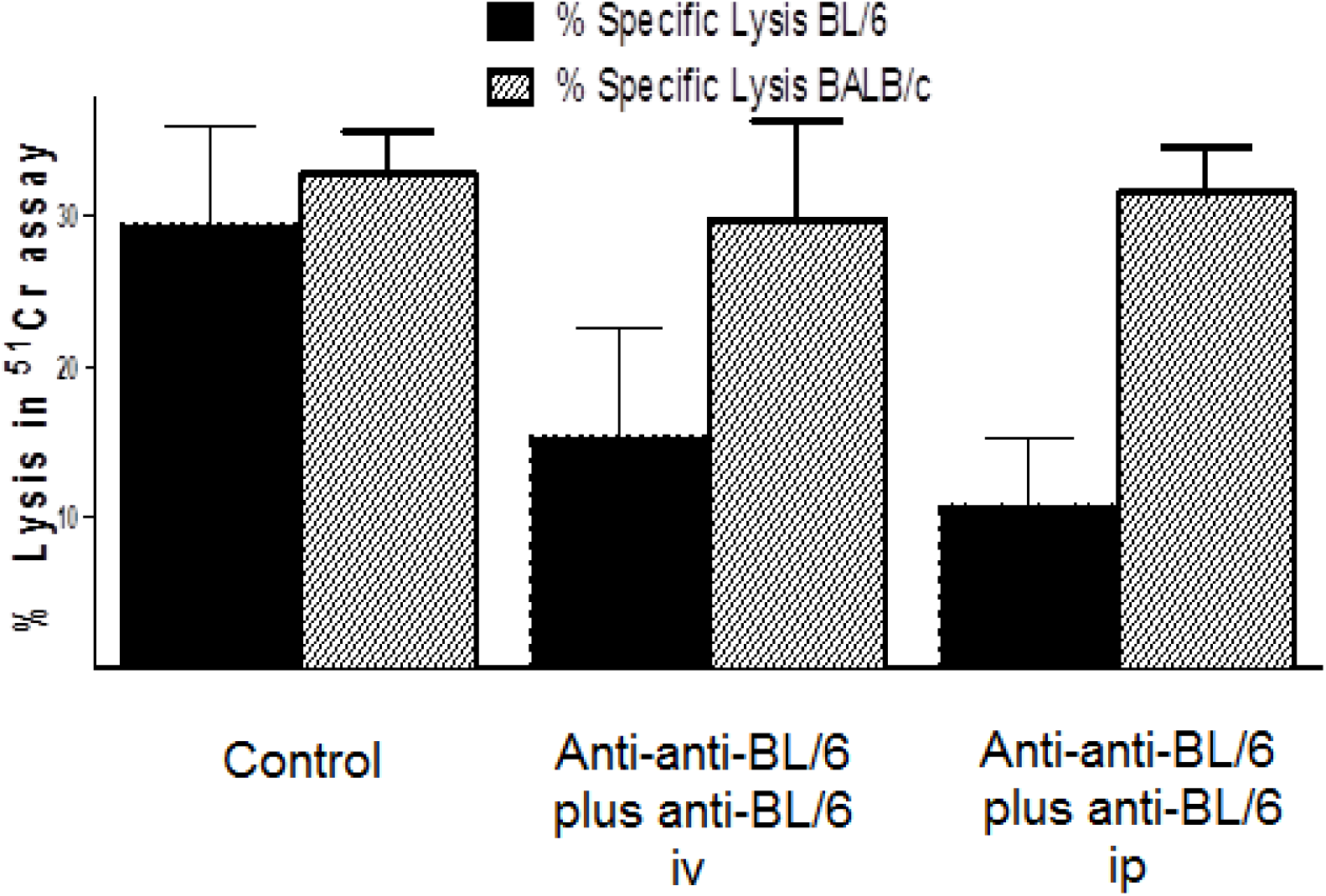
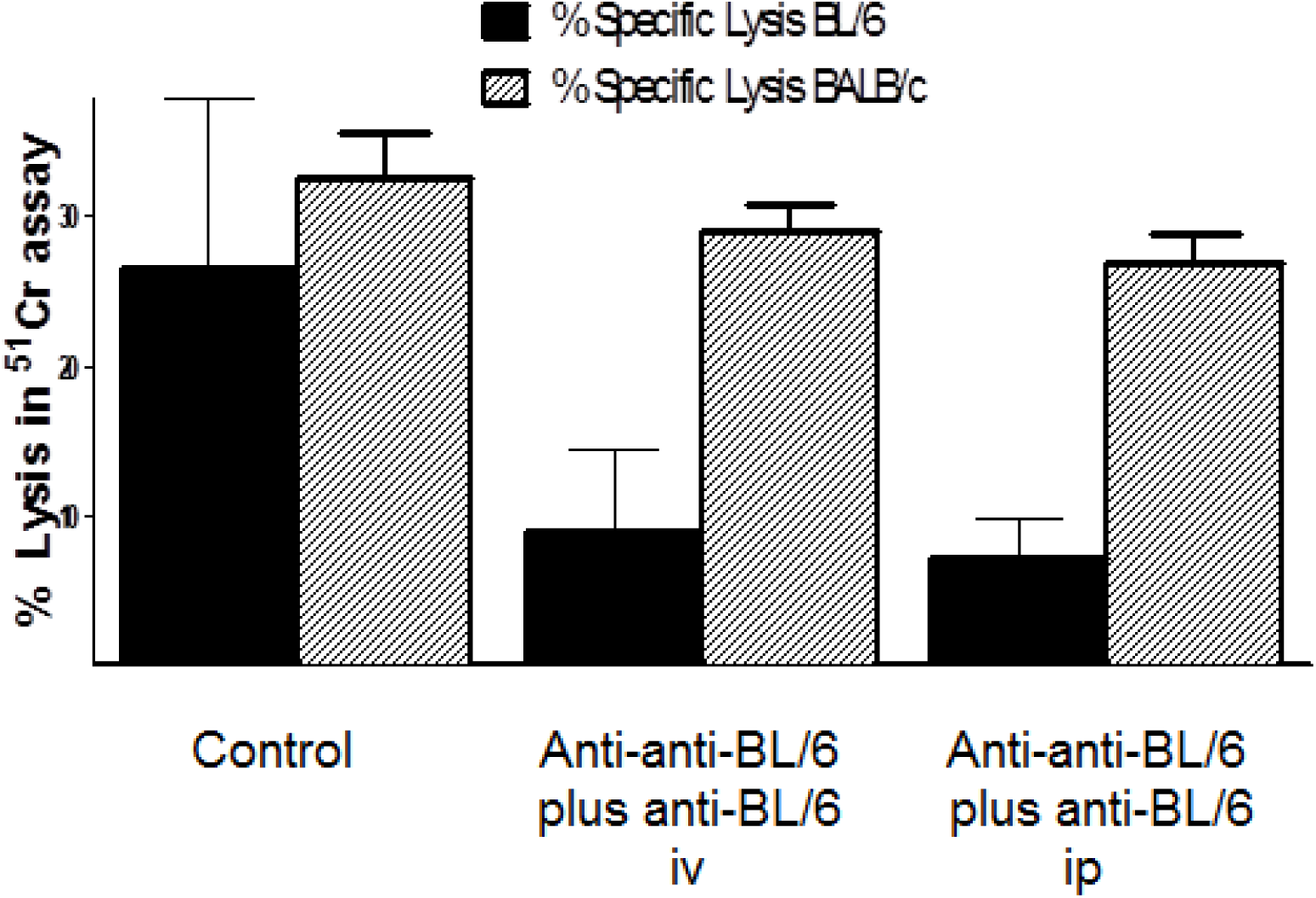
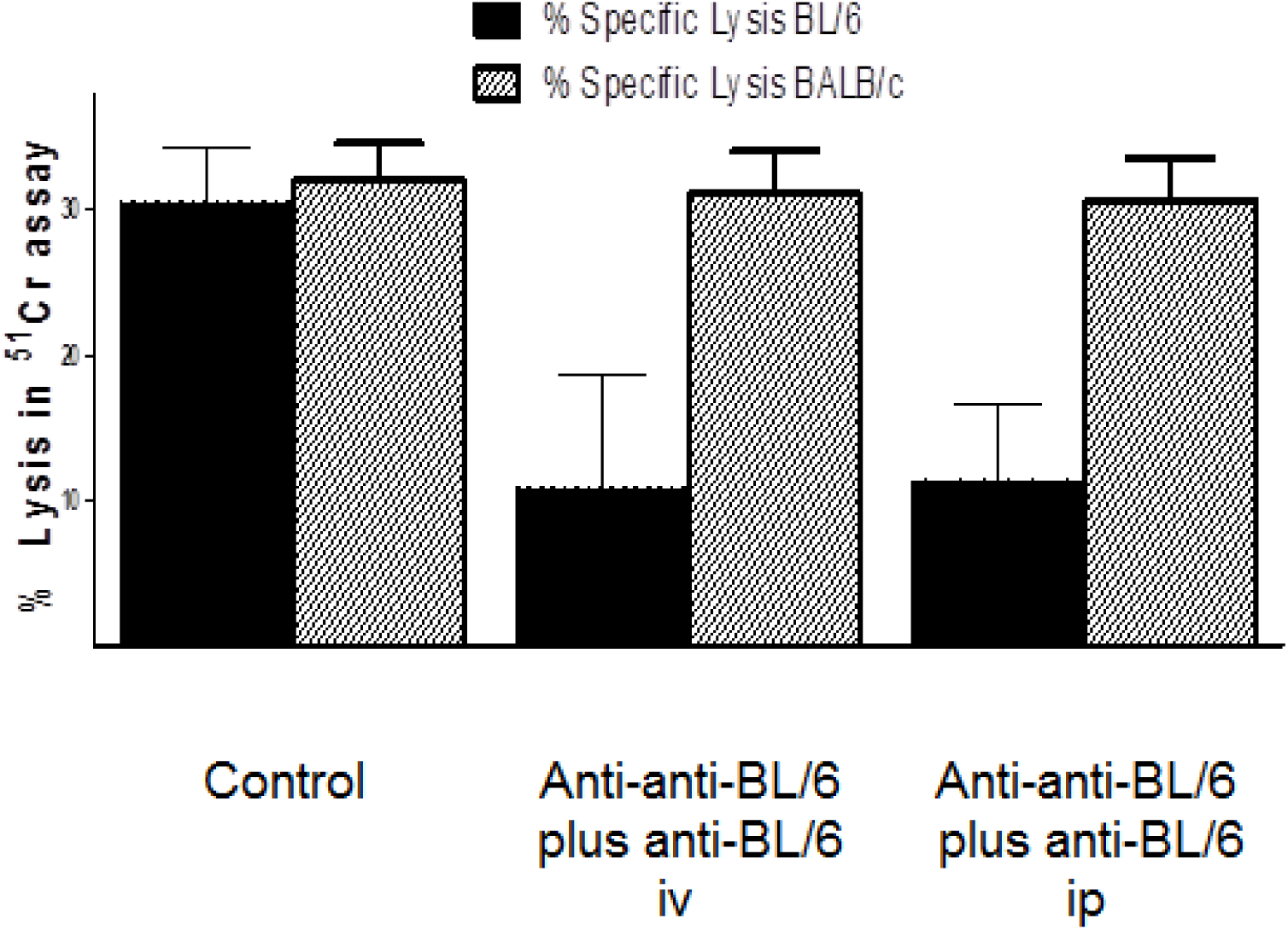
Suppression of anti-BL/6 cytotoxic T cell generation by anti-BL/6 plus anti-anti-BL/6 antibodies. Figures 3A to 3E. Anti-BL/6 CTL response of lymphocytes from C3H mice that had been treated with weekly infusions of 1μg anti-BL/6 IgG plus 1μg anti-anti-BL/6 IgG given either intravenously or intraperitoneal. Figure 3A shows the CTL response after two infusions, Figure 3B after three infusions, Figure 3C after four infusions, Figure 3D after five infusions and Figure 3E after six infusions.

The anti-BL/6 CTL response of lymphocytes from C3H mice that had been treated with weekly infusions of anti-BL/6 IgG plus anti-anti-BL/6 IgG given either intravenously or intraperitoneal is shown in Figures 3A to 3E. Anti-BALB/c CTL is the control. Figure 3A shows the CTL response after two infusions, Figure 3B after three infusions, Figure 3C after four infusions, Figure 3D after five infusions and Figure 3E after six infusions. The mice were sacrificed at day 42 and tested for the induction of BALB/c-specific and BL/6-specific CTL.

In a second experiment with different controls, groups of 10 C3H mice received saline (Group 1), 10ug anti-BL/6 IgG (Group 2), 10ug anti-anti-BL/6 IgG (Group 3) or 10ug anti-BL/6 plus 10ug anti-anti-BL/6 IgG (Group 4) on days 0, 7, 14, 21, and 28. We sacrificed the mice at day 42 and tested for the induction of BL/6-specific and BALB/c-specific CTL. The results are shown in Figure 3F. This experiment shows that the combination of anti-BL/6 plus anti-anti-BL/6 IgG synergistically causes a significantly reduced CTL response compared to each of the other groups (P < 0.01 by ANOVA).

**Figure 3F.**
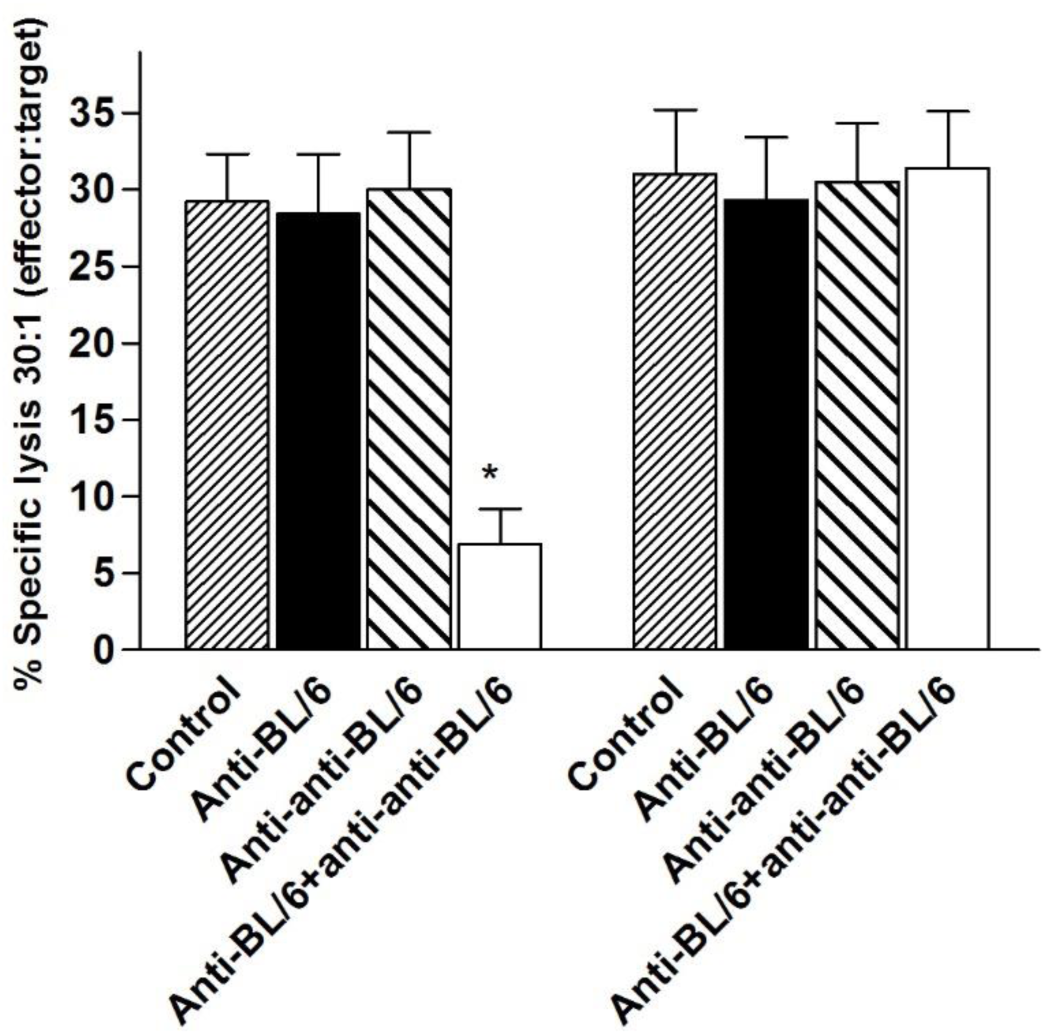
Groups of C3H mice were given 10μg C3H anti-BL/6 plus 10μg BL/6 anti-anti-BL/6 (Group 1), or 10μg BL/6 anti-anti-BL/6 (Group 2), or 10μg C3H anti-BL/6 (Group 3), or phosphate buffered saline at neutral pH on days 0, 7, 14, 21 and 28. On day 42 they were sacrificed and spleen cells were cultured with BL/6 or BALB/c stimulator cells and BL/6-specific and BALB/c-specific cytotoxicity respectively was measured. 20 mice per Group. * denotes P < 0.01 by ANOVA.

Together these experiments show that anti-BL/6 plus anti-anti-BL/6 synergistically induce specific tolerance in C3H mice. More generally, we have experimental results together with a plausible theoretical mechanism (Figure 2A), so we expect that stimulation of an immune system by antigen-specific (for example anti-B) plus complementary antiidiotypic antibodies (anti-anti-B) can generally induce a new stable steady state in which the immune response to B is specifically suppressed.

### Synergy of antigen-specific and antiidiotypic antibodies in preventing inflammatory bowel disease in mice

Inflammatory bowel disease (IBD) is a category of autoimmune diseases that includes colitis and Crohn’s disease. BL/6 mice fed dextran sodium sulphate (DSS) develop IBD. We tested the efficacy of prevention of IBD by antigen-specific plus antiidiotypic antibodies in an experiment with three groups of eight BL/6 mice. The first group was given a normal diet, the second was given a diet that included DSS and high fat, and the third was given DSS and high fat plus two infusions of antigen-specific (BL/6 anti-C3H) plus antiidiotypic (C3H anti-anti-C3H) antibodies for two weeks at days −14 and −7 prior to commencement of the diet of the DSS plus high fat diet at day 0. The mice were successfully treated by the combination of antibodies as measured in three assays, namely significant inhibition of the production of mRNA for seven of nine inflammatory cytokines (Figure 4), reduction in the change of colon length (Figure 5) and reduction of weight loss (Figure 6) (P < 0.05 by ANOVA). We attribute this therapeutic effect to changing the phenotype of the treated mice to have similarity to that of C3H mice, while not losing tolerance to the BL/6 phenotype, meaning self-tolerance to BL/6 is not lost. The treated BL/6 mice have the benefit of tolerance to both BL/6 and C3H, while the ability to respond to other antigens is fully retained. In a control experiment (not shown) the mice were treated with only C3H anti-anti-C3H antibodies. The suppression of inflammation was then not observed. We conclude that treatment with antigen-specific plus antiidiotypic antibodies synergistically prevents inflammatory bowel disease.

**Figure 4.**
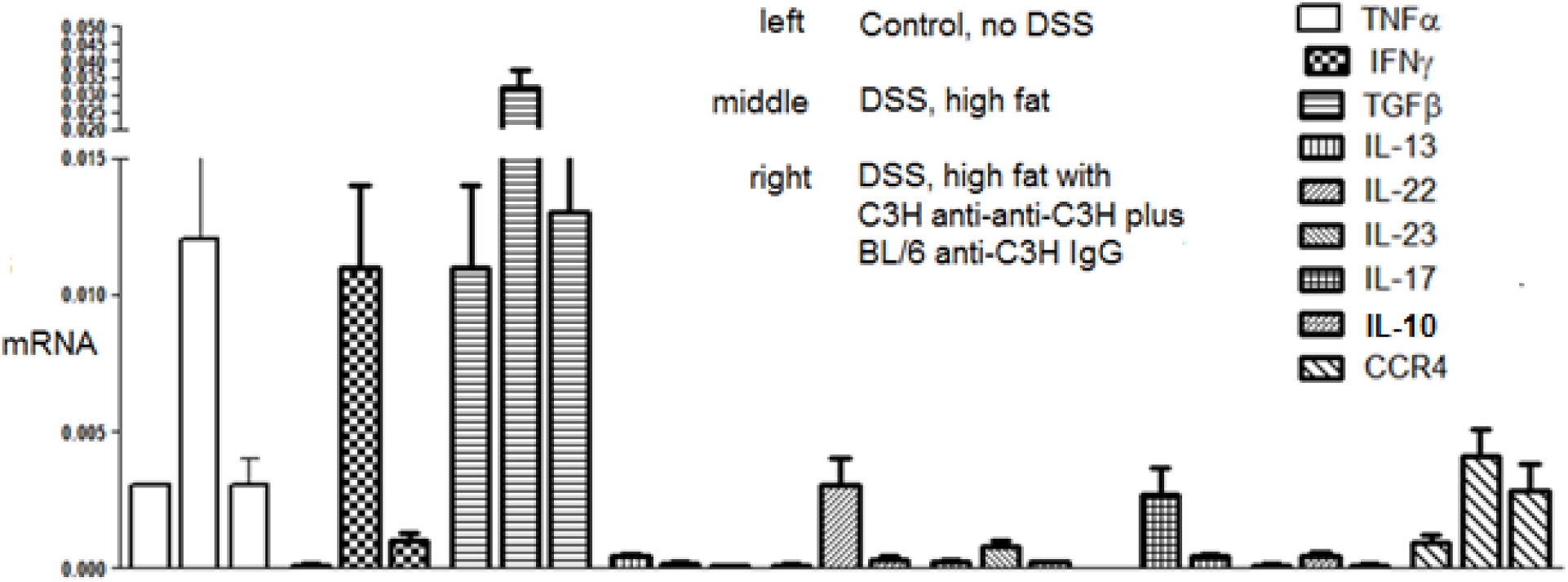
Induction of cytokine mRNA production in BL/6 mice caused by dextran sodium sulphate (DSS) and high fat diet, and inhibition of the cytokine mRNA production caused by C3H anti-anti-C3H plus BL/6 anti-C3H IgG antibodies. The mice were given 10μg of each of the antibodies intravenously on days −14 and −7 prior to the commencement of DSS and high fat diet on day 0. Eight mice per group.

**Figure 5.**
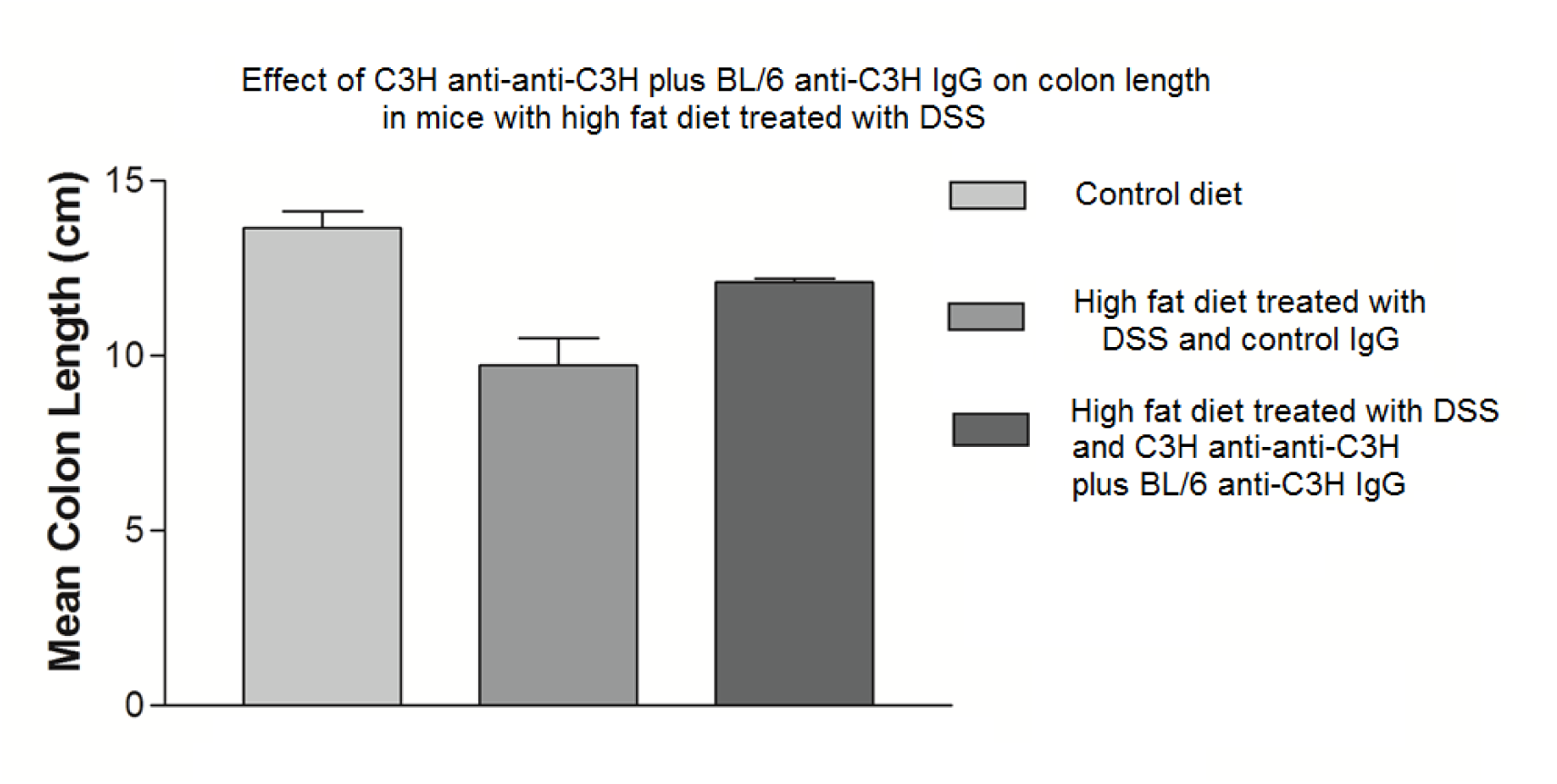
Change in colon length caused by dextran sodium sulphate and high fat diet, and decrease in the change caused by C3H anti-anti-C3H antibodies plus BL/6 anti-C3H IgG antibodies.

**Figure 6.**
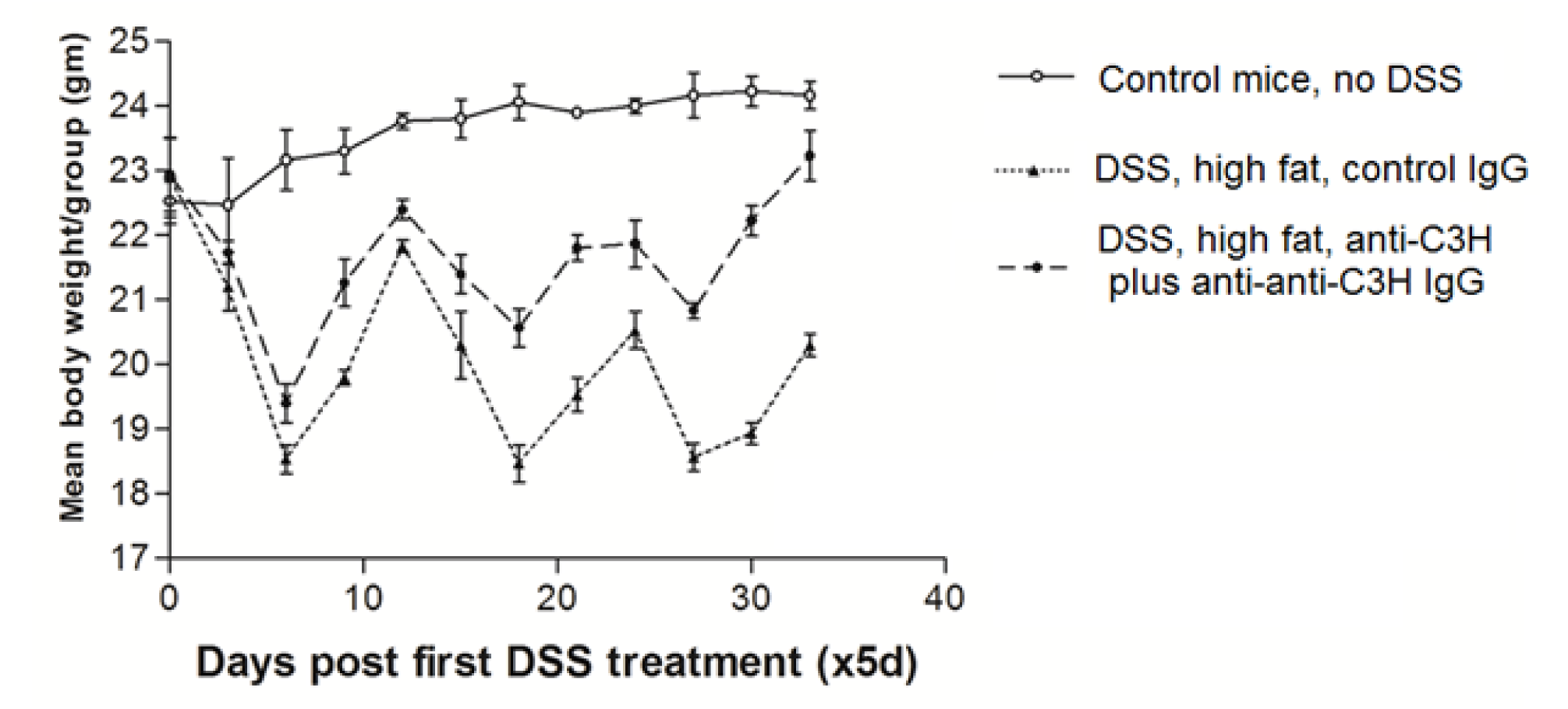
Weight loss caused by DSS and high fat diet, and reduction in the weight loss caused by C3H anti-anti-C3H antibodies plus BL/6 anti-C3H IgG antibodies.

### Synergy of antigen-specific and antiidiotypic antibodies in prevention of breast cancer in a mouse model

EMT6 is a transplantable breast cancer tumour (Gorczynski et al., 2013). 6 BALB/c mice/group received 5×10^5^ EMT6 cells 14d after 2 intravenous injections of C3H anti-BL/6 and/or BL/6 anti-anti-BL/6 IgG (10μg/mouse). Injections were continued at 7 day intervals thereafter until final sacrifice. 14 days after EMT6 injection mice were anesthetized and tumors resected and weighed. Mice were returned to their cages for a further 16 days when all were sacrificed and lung nodules evaluated. The results for tumour size and the number of metastases in the lungs are shown in Figure 7. * indicates P < 0.05 compared with the other groups. The tumour size and the number of metastases were significantly less in the mice that received C3H anti-BL/6 plus BL/6 anti-anti-BL/6 IgG.

**Figure 7.**
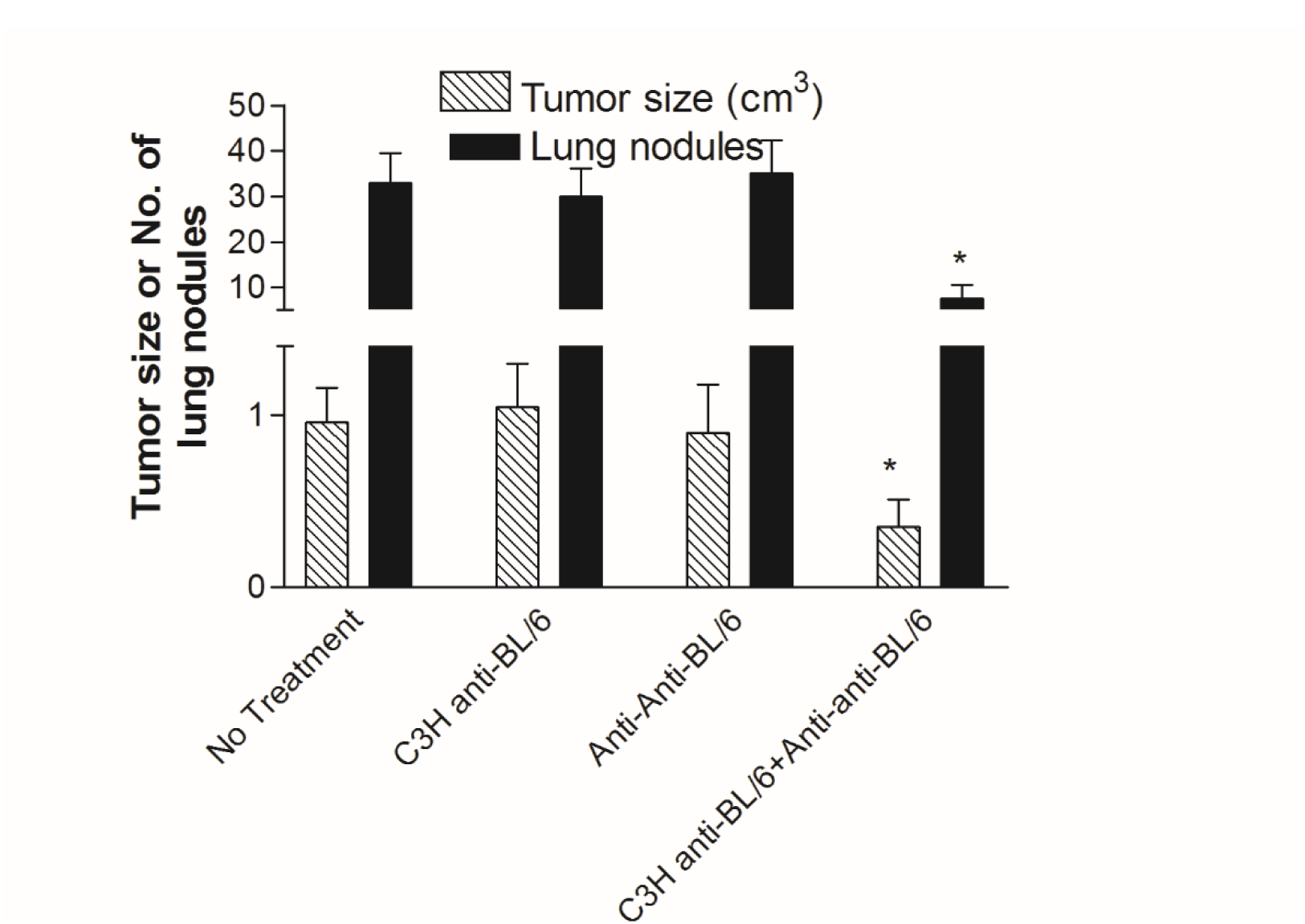
Decrease in tumour growth and decrease in metastases in BALB/c mice given anti-foreign (C3H anti-BL/6) plus complementary antiidiotypic (BL/6 anti-anti-BL/6) antibodies and the transplantable breast cancer EMT6. Six mice per group received 5x10^5^ EMT6 cells on day 0 after 2 intravenous injections of anti-BL/6 and/or anti-anti-BL/6 Ig (10μg/mouse) on days −14 and day −7. Injections were continued at 7d intervals thereafter until final sacrifice. 14 days after EMT6 injection the mice were anesthetized and tumors resected and weighed. The mice were returned to their cages for a further 16d when all were sacrificed and lung nodules evaluated. * indicates p<0.05 compared with all other groups.

Cytokine data for this experiment is shown in Figure 8. Lymph nodes draining the tumor were harvested at sacrifice, and 1×10^6^ cells were cultured in duplicate in 1ml medium with either 1×10^6^ irradiated (2500Rads) BL/6 splenocytes or 1×10^5^ irradiated EMT6 tumor cells. Supernatants were harvested at 48hr and assayed in commercial ELISAs (BioLegend) for the cytokines shown. * indicates p < 0.05 compared with all other groups. The results for stimulation of BL/6 spleen cells (H-2^b^) and of EMT6 (H-2^d^) are the same. IL-1β and IL-6 are significantly down-regulated, while TGFβ is significantly up-regulated. IL-1β and IL-6 are pro-inflammatory cytokines while TGFβ regulates inflammation. There was no change in the level of TNFα. Hence for three of the four cytokines there is a significant therapeutic effect resulting from the infusions of the antigen-specific plus antiidiotypic antibodies.

**Figure 8.**
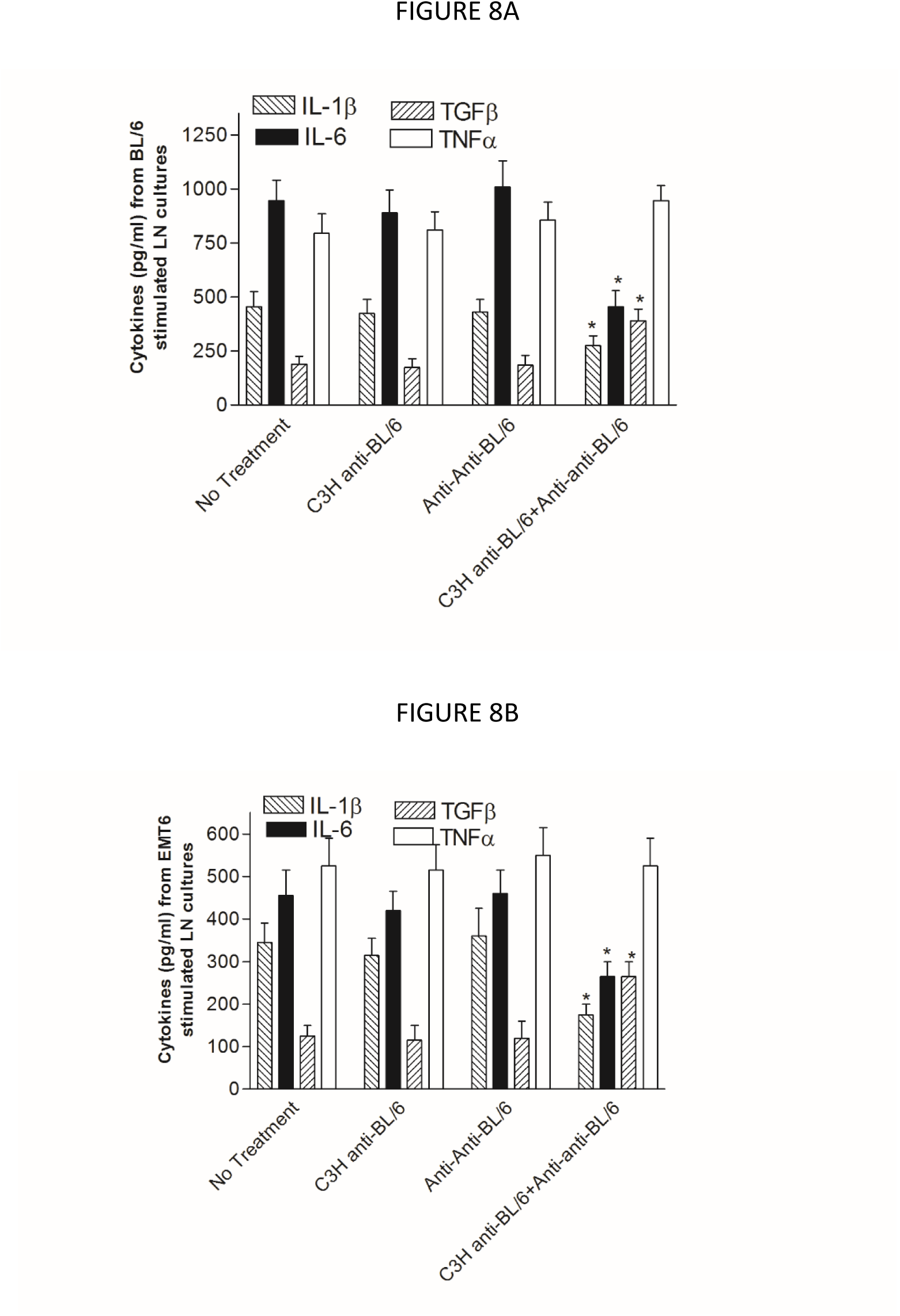
Cytokine production for lymphocytes from the experiment of Figure 7. Lymph nodes draining the tumor were harvested at sacrifice, and 1×10^6^ cells cultured in duplicate in 1ml medium with either 1×10^6^ irradiated (2500Rads) BL/6 splenocytes (Figure 8A) or 1×10^5^ irradiated EMT6 tumor cells (Figure 8B). Supernatants were harvested at 48hr and assayed in commercial ELISAs (BioLegend) for the cytokines shown. * indicates p<0.05 compared with all other groups by ANOVA.

## Discussion

The symmetrical immune network theory has been developed over a period of more than forty years, and accounts for many aspects of the adaptive immune system (Hoffmann et al., 1986; Hoffmann, 1994; Hoffmann, 2008; Leung and Hoffmann, 2014). An important aspect of the theory is that it is based on the controversial presumed existence of specific T cell factors (tabs) (Hoffmann, 1991; Melchers, 1990; Vaz and Coutinho, 1991). We recently confirmed (Hoffmann and Gorczynski, 2017) that tabs do indeed exist by reproducing a rigorous experiment published in 1975 by Takemori and Tada (1975).

We find that normal IgG immune responses in a vertebrate against an antigen consist of two components, namely antigen-specific and second symmetry antiidiotypic antibodies. The second symmetry antiidiotypic antibodies in an immune response of a vertebrate A to skin grafts from a vertebrate B (A anti-anti-A) bind to antigen-specific antibodies present in a B anti-A immune response. The IgG of a vertebrate A immune to the protein antigen tetanus toxoid likewise includes antiidiotypic antibodies (A anti-anti-A) that are specific for antigen-specific antibodies (B anti-A) present in a B anti-A immune response.

Antigen-specific plus second symmetry antiidiotypic antibodies synergistically induce tolerance. The proposed mechanism for this is that there is co-selection of antigen-specific and complementary antiidiotypic T cells. This combination of antibodies is also effective in down-regulating inflammation in a murine model of inflammatory bowel disease as measured in each of three assays. This combination is also effective in significantly down-regulating tumour growth and preventing metastases in the EMT6 mouse breast cancer model. In this case two of three pro-inflammatory cytokines measured are down-regulated, and an anti-inflammatory cytokine is upregulated.

Our experiments indicate that the immune systems of two healthy vertebrates called A and B can be combined with the immune system of a third healthy vertebrate C to make C’s immune system resistant to degenerative diseases. A, B and C may be three strains of mice, and in the case of the application of the immunotherapy to humans, A and B may be mouse strains and C may be a healthy human. Anti-B antibodies may be obtained from immunization of A with B tissue, and anti-anti-B antibodies may be obtained from immunization of B with A tissue. C is then treated with a combination of anti-B and anti-anti-B antibodies. This is the basis for a preventive immunotherapy comprising infusions with two mutually specific antibodies.

Why has evolution not led to normal immune systems being similarly resistant to degenerative diseases? To address this question, we first consider that, according to the symmetrical immune network theory normal immune systems include self antigens C, lymphocytes that are anti-C and are co-selected with anti-anti-C lymphocytes, and lymphocytes that are anti-anti-anti-C and are also co-selected with anti-anti-C lymphocytes, as shown in Figure 9 (Hoffmann, 2008, chapter 17). The antigens C include especially MHC class II antigens that stimulate anti-C Th1 lymphocytes and Ts1 lymphocytes. The anti-anti-C lymphocytes include Th2, Ts2 and B2 lymphocytes, whereby B2 lymphocytes are B cells that secrete IgG. The anti-anti-anti-C lymphocytes include Ts3 and B1 lymphocytes, whereby B1 lymphocytes secret IgM. There is therefore a principle “shape space axis” comprising C, anti-C, anti-anti-C and anti-anti-anti-C. Each of the members in this sequence has shapes that are complementary to the next members in the sequence. We can also depict the shape space axis in a way such that neighboring members are similar to each other, and members with complementary shapes are on opposite sides of the diagram. This is shown in Figure 10A, in which the axis is defined by anti-C and anti-anti-C. C is complementary to anti-C, so it is on the same side as anti-anti-C, and anti-anti-anti-C is complementary to anti-anti-C so it is on the same side as anti-C. The system is stabilized by anti-C being stimulated by anti-anti-C and vice versa. At any given time, there is a mixture of anti-C and anti-anti-C factors on the A cell surface, and we could expect random fluctuations in which of the two (anti-C or anti-anti-C) is present to a greater degree. Such fluctuations would result in fluctuations in the number of anti-C and anti-anti-C lymphocytes. There may then be a random walk in one dimension for the relative amounts of anti-C and anti-anti-C lymphocytes. At some stage this random walk may lead to the two populations no longer being mutually stabilizing. This idea is supported by the fact that autoimmune mice make anti-anti-C antibodies (Kion and Hoffmann, 1991), which could be due to T cells no longer dominating the anti-anti-C region of shape space. In addition, we have found that old BL/6 mice make anti-anti-BL/6 (or “anti-anti-C”) antibodies (Al Fahim and Hoffmann, 1996).

**Figure 9.**
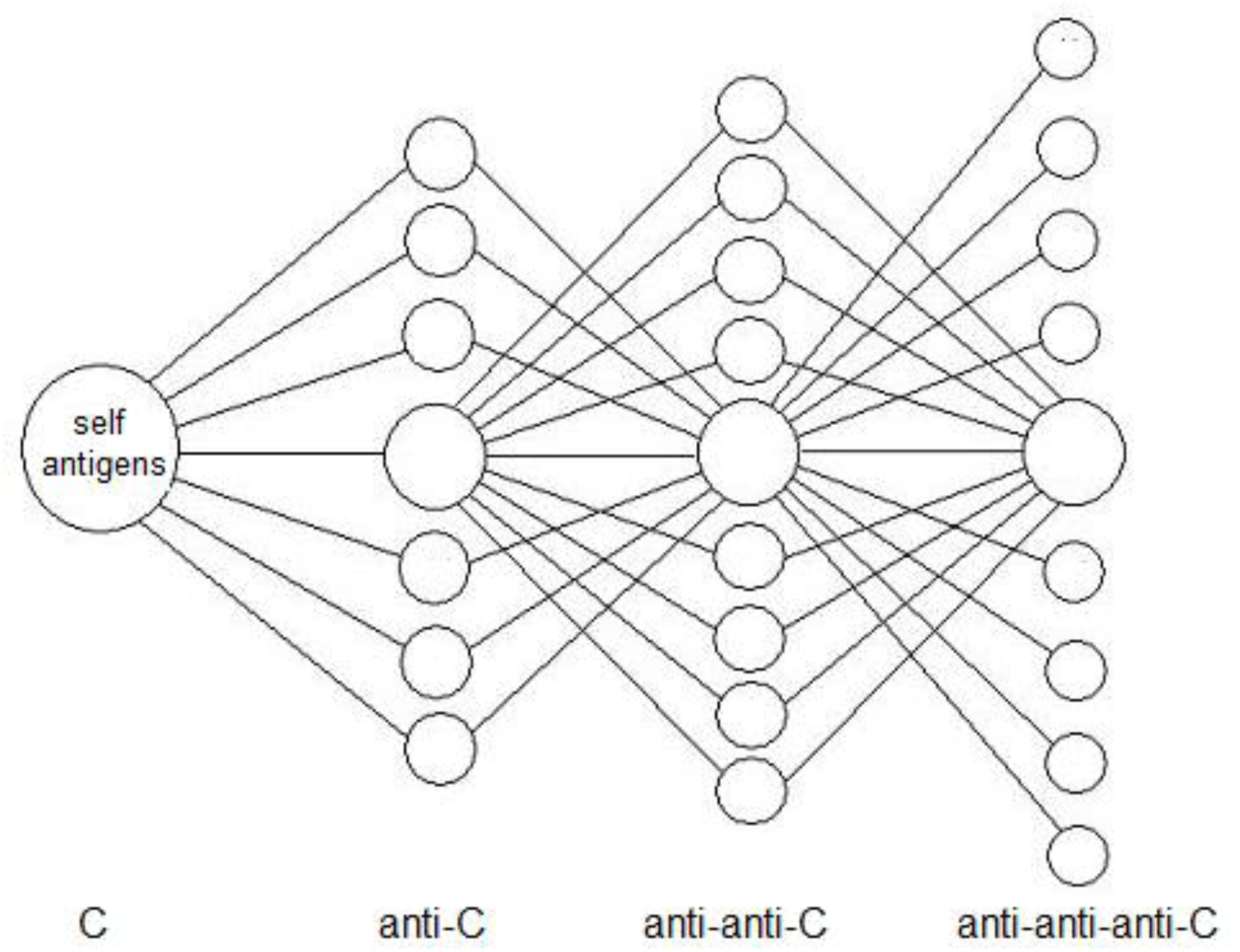
A model that is part of the symmetrical immune network theory, adapted from Figure 17-3 of Hoffmann, 2008. Self antigens of a vertebrate C stimulate anti-C (including Th1 and Ts1 lymphocytes), which are co-selected with anti-anti-C (including Th2, Ts2 and B2 lymphocytes), witch in turn are co-selected with anti-anti-anti-C (including Ts3 and B1 lymphocytes). B2 cells are IgG-secreting B lymphocytes, and B1 cells are IgM-secreting B lymphocytes.

**Figure 10.**
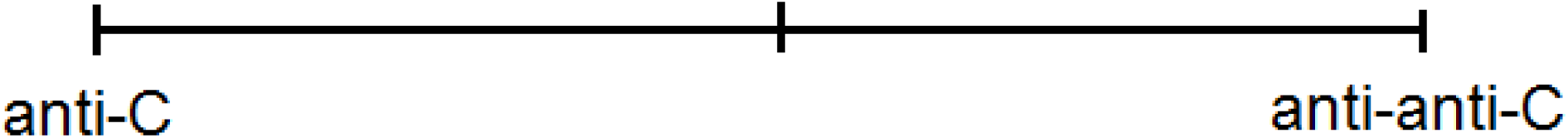
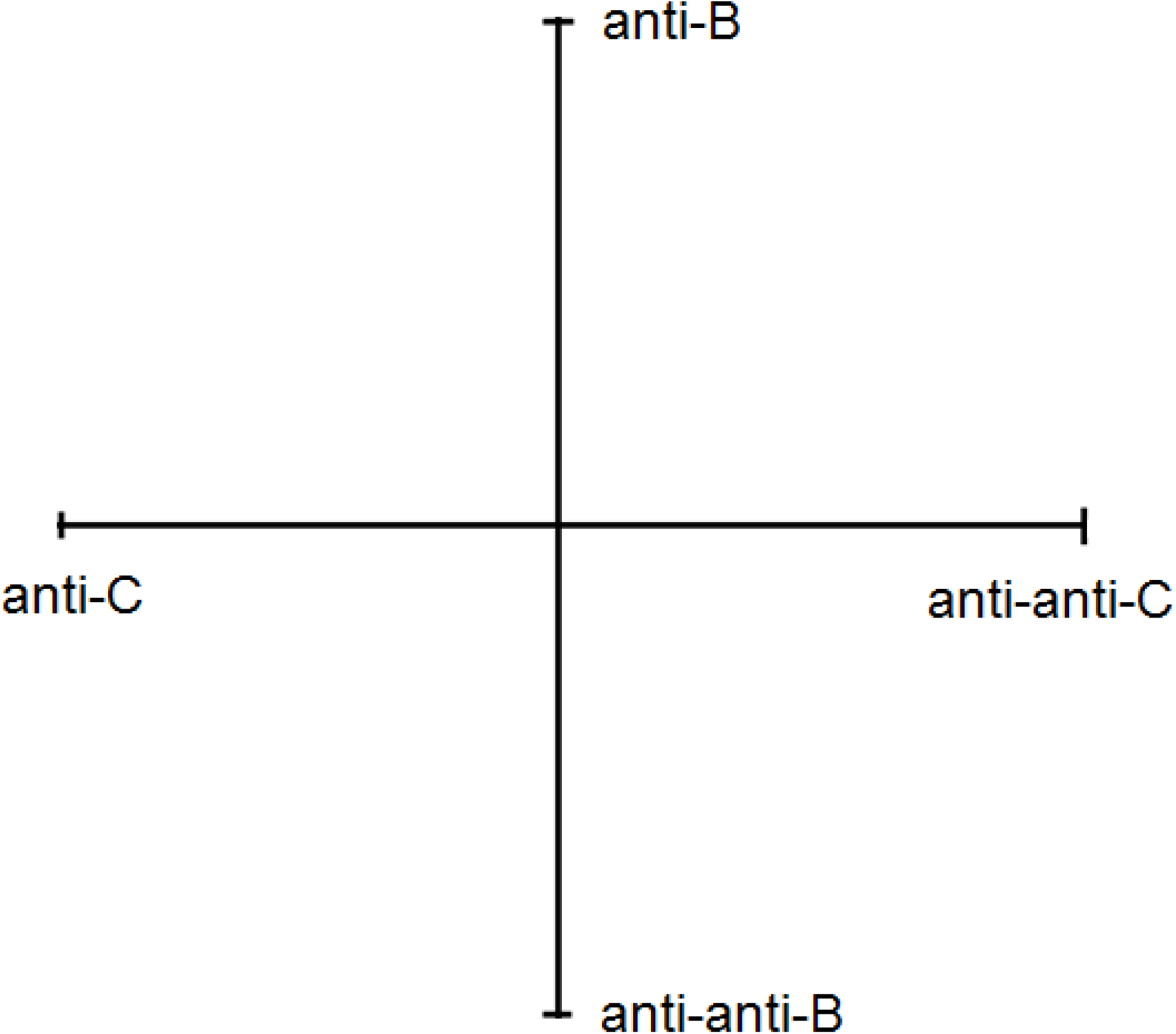
One-dimensional and two-dimensional shape space models. Figure 10A. A onedimensional shape space model for the vertebrate C in which complementary shapes map to opposite sides of the origin. Figure 10B. A two-dimensional shape space model results when the vertebrate C is treated with anti-B plus anti-anti-B antibodies. Anti-anti-C and anti-anti-B lymphocytes are co-selected with anti-C and anti-B lymphocytes respectively.

When we treat C with anti-B plus anti-anti-B antibodies we induce a new stable steady state with elevated levels of anti-anti-B and anti-B T cells. This amounts to creating a second shape space axis for the T cells of C that is plausibly orthogonally to the anti-C/anti-anti-C shape space axis because neither anti-B nor anti-anti-B has any a priori relation to anti-C or anti-anti-C. See Figure 10B. In this case fluctuations in all of anti-C, anti-anti-C, anti-B and anti-anti-B specific T cell factors would mean there is a random walk in two dimensions rather than one dimension. An immune system with a single shape space axis as shown in Figure 10A may then be intrinsically less stable that a system with a second shape space axis as shown in Figure 10B. In the case of Figure 10A the steady state involves primarily just anti-C and anti-anti-C T cell factors on the A cell surface, while there are four specificities of specific T cell factors for a system with two shape space axes as shown in Figure 10B. This difference in the number of shape space dimensions may be the basis for an immune system constructed using three immune systems (A, B and C), and having two shape space axes, being intrinsically stronger than an immune system based on a single shape space axis.

We conclude that second symmetry antiidiotypes and induced co-selection is leading to a possible preventive immunotherapy, that does not involve the production of antibodies and can reasonably be called a new class of vaccine. Combinations of antigen-specific antibodies and second symmetry antiidiotypic antibodies as described here comprise an immunotherapy for the prevention of inflammatory bowel disease and the prevention of breast cancer. The same therapy may prove to be effective in preventing also other autoimmune diseases and cancers.

## MATERIALS AND METHODS

### Skin grafting

C3H or BL/6 mice received allogeneic or syngeneic skin grafts transplanted to the flank as described previously. Graft survival was monitored daily by an observer blinded to any previous treatment of the graft recipients.

In some cases grafted mice were used as sources of anti-graft specific Ig (or anti-anti-self Ig). In these instances mice were grafted twice (at 21d intervals) with donor skin (same donor haplotype) and blood obtained by cardiac puncture 10d after the second graft. Serum was obtained by centrifugation (5000g at 4°C for 20 min), heat inactivated, aliquoted (0.3ml aliquots) and stored at −80°C. Where serum was absorbed (anti-anti-donor Ig), aliquots (0.3ml) were absorbed 3x for 60min at room temperature with a fresh pellet of 3×10^8^ spleen/thymus, prepared from the described donors. Following 3 serial absorptions, depletion of cytotoxic activity in serum was confirmed using spleen cell blasts and serial dilutions of antibody with rabbit complement (1:10 final dilution) incubated at 37°C for 60min before addition of trypan blue to assess cell death. Routinely titres dropped from 50% lysis at ˜1:2000 to ˜1:2 following this absorption.

In other studies, mice were pretreated before transplantation with A anti-B sera and/or anti-anti-self Ig. In these cases, animals received weekly injections (intravenous, iv, and/or intraperitoneal, ip) of serum IG at the concentration described, diluted in 0. 3ml PBS. In addition, and again as described in the text, in some instances mice received injections of the broadly neutralizing anti-HIV antibody b12. This was purchased from Polymun Scientific Immunbiologische Forschung GmbH, Klosterneuburg, Austria.

### Mixed leukocyte culture assays (MLCs) and ^51^Cr lysis assays for CTL

Graft recipients were sacrificed at the times described in the text/Figure legends, and single cell splenocyte suspensions prepared for individual mice. 6×10^6^ responder cells were incubated in triplicate with 3×10^6^ irradiated spleen stimulator cells in flat bottomed culture wells (24-well culture plates) in 3.0 ml MEM supplemented with 10% Fetal Calf serum and 10^−6^M 2-mercaptoethanol (F10). In some cultures, an aliquot (20001) of medium was removed at 40hrs to assess cytokine production, using commercial ELISA kits (BioLegend, USA). After 5d cells were harvested, washed, and used at different effector:target ratios (from 50:1 to 5:1) in triplicate for lysis over 5hrs of 5×10^3^ ^51^Cr-labeled 72hr ConA spleen cell blasts homologous with the cells used as stimulator in MLCS. Data shown in Figures represent mean lysis at a 30:1 E:T ratio.

### IBD after IV Ig treatment

#### Mice

C57BL/6 female mice, with high susceptibility to DSS-induced colitis, purchased from the Jackson laboratories (Bar Harbor, ME). were used throughout. All mice were housed five per cage under specific pathogen-free conditions, allowed standard diet (or high fat) and water ad libitum, and used at 6-8 wk of age. Where shown mice received 10g BL/6 anti-C3H (ip) and l0g C3H anti-BL/6 absorbed with BL/6 (C3H anti-anti-C3H) iv weekly beginning 14d before the first DSS treatment.

Animal experimentation was performed following guidelines of an accredited animal care committee (protocol no. AUP.1.5). Humane endpoints were used in all studies, with mice monitored daily. Animals were euthanized (overdose with pentobarbital) when they were exhibiting signs of distress (weight loss≥25%; hunched posture; diaorrhea; loss of active movements)-no mortality was seen in this chronic IBD protocol. Animals with diaorrhea (but weight loss <25%) received daily ip injections of saline (1ml x3 at 8hr intervals) to avoid dehydration.

#### Induction of colitis

Chronic DSS colitis was induced by giving mice distilled drinking water containing 3% (wt/vol) DSS (m.w. = 40 kDa; ICN Biochemicals, Aurora, OH) for 5 days followed by 7 days of normal drinking water for a total of 3 cycles. Body weight was measured three times/week throughout the experiment. No significant mortality was seen in any groups in the chronic colitis model. As shown in Figures maximum weight loss was ˜25% with all mice recovering weight loss within 7-10 days post cessation of DSS exposure.

#### RNA isolation and real-time RT-PCR

Total RNA was isolated from colonic tissue using TRIzol reagent (Invitrogen Canada, Burlington, ON). Total RNA (1 μg) was treated with DNase I and then reverse transcribed using High Capacity cDNA Reverse Transcription Kit (Applied Biosystems, Foster city, CA) following the manufacturer’s instruction. First strain cDNA was diluted 1:20 and used for quantitative PCR on an ABI 7900HT Sequence Detection system with SYBR green master mix (Applied Biosystems). Messenger RNA levels were normalized to a composite of GAPDH and HPRT expression levels. Experiments were repeated three times with three independent cDNA syntheses. The primers used for real-time RT-PCR are shown below.

Primers used for real-time PCR

IFN-γ forward: 5'-TATT GCCAAGTTT GAGGTCAACA-3′ reverse: 5′-GCTGGATTCCGGCAACAG-3′
TNF-α forward: 5'-AGACCCTCACACTCAGATCATCTTC-3′ reverse: 5′-CCACTTGGTGGTTTGCTACGA-3′
IL-1β forward: 5′-TCGTGCTGTCGGACCCAT AT-3′ reverse: 5′-GGTTCTCCTTGTACAAAGCTCATG-3′
IL-4 forward: 5′-TCAT CGGCATTTTGAACGAG-3′ reverse: 5′-TTTG GCACATCCAT CTCCG-3′
IL-6 forward: 5′-CTCTGGGAAATCGTGGAAATG-3′ reverse: 5′-CAGATTGTTTTCT GCAAGT GCAT-3′
IL-10 forward: 5′-AAGGCAGTGGAGCAGGTGAA-3′ reverse: 5′-TTCTATGCAGTTGATGAAGATGTCAA-3′
IL-12 forward: 5′-CCCAAGGT CAGCGTT CCA-3′ reverse: 5′-GGCAAGGGTGGCCAAAA-3′
IL-17 forward: 5′-CTCAGACTACCTCAACCGTTCCA-3′ reverse: 5′-CCAGATCACAGAGGGATATCTATCAG-3′
IL-22 forward: 5′-GTGCCTTTCCTGACCAAA-3′ reverse: 5′-TCTCCTTCAGCCTTCTGA-3′
IL-23 forward: 5′-GACAACAGCCAGTTCT GCTT-3′ reverse: 5′-AGGGAGGTGTGAAGTTGCTC-3′
IL-23R forward: 5′-AATTTGACGCCAATTTCACA-3′ reverse: 5′-ACCAGTTT CTT GACAT CGCA-3′
TGF-β forward: 5′-CGAAGCGGACTACTATGCTAAAGA-3′ reverse: 5′-GTTTTCTCATAGATGGCGTTGTTG-3′
Foxp3 forward: 5′-AGTCTGCAAGTGGCCTGGTT-3′ reverse: 5′-GGGCCTTGCCTTTCTCATC-3′
CCR4 forward: 5′-AGACTGT CCTCAGGAT CACTTTCA-3′ reverse: 5′-CCGGGTACCAGCAGGAGAA-3′
CCL-17 forward: 5′-ATGCCATCGTGTTTCTGACTGT-3′ reverse: 5′-GCCTTGGGTTTTTCACCAAT C-3′
CCL-22 forward: 5′-AAGCCTGGCGTTGTTTTGAT-3′ reverse: 5′-TCCCTAGGACAGTTTATGGAGTAGCT-3′ reverse: 5′-AAGCCGAGTTCAGCAAAGTT-3′

#### Groups

1. 5 mice were controls given a normal diet (details)
2. 5 mice were given DSS
3. 5 mice were given DSS together with anti-anti-C3H and anti-C3H IgG antibodies

#### Colonic explant culture and multi-analyte Elisarray

Colons were removed, washed three times with cold PBS containing 100 IU penicillin and 100 μg/ml streptomycin, and opened longitudinally. The proximal and distal parts of the colon were cut into strips 3 mm in length and cultured in 500 μl of supplemented RPMI 1640 culture medium at 37 °C with 5% CO2 humidified air for 24 h following which supernatants were collected and particulate material removed by centrifugation for 10 min at 1000 x g. The supernatants were subsequently used for cytokine and chemokine analysis using a Multi-analyte Elisarray kit (Qiagen, Mississauga, ON). Twelve proinflammatory cytokines and chemokines were examined in the supernatants, including IL-1β, IL-4, IL-6, IL-10, IL-12, IL17A, IFN-γ, TNF-α, TGFβ, MCP1, MIP-1α and MIP-1β. Capture antibodies for the 12 cytokines and chemokines were coated on one Elisarray microplate and 50 μl of samples were added to the wells of the plate. After 2 h incubation and exhaustive washing to remove unbound proteins, 100 μl of biotinylated detection antibodies were added. Thereafter, an avidin-horseradish peroxidase conjugate was added after 1 h incubation. After further washing substrate solution was added. A stop solution was added after 30 min and the absorbance at 450 nm was read. The manufacturer’s computer software, which includes relevant cytokine controls, was used to express the cytokine “signals” in different samples relative to one control group (nominally scored as =1).

#### Weight loss

The weights of the same groups of mice were monitored daily, and the mean and standard deviation were plotted as a function of time.

